# AMRomics: a scalable workflow to analyze large microbial genome collection

**DOI:** 10.1101/2024.04.02.587817

**Authors:** Duc Quang Le, Tam Thi Nguyen, Canh Hao Nguyen, Tho Huu Ho, Nam S. Vo, Trang Nguyen, Hoang Anh Nguyen, Minh Duc Cao, Son Hoang Nguyen

## Abstract

Whole genome analysis for microbial genomics is critical to studying and monitoring antimicrobial resistance strains. The exponential growth of microbial sequencing data necessitates a fast and scalable computational pipeline to generate the desired outputs in a timely and cost-effective manner. Recent methods have been implemented to integrate individual genomes into large collections of specific bacterial populations and are widely employed for systematic genomic surveillance. However, they do not scale well when the population expands and turnaround time remains the main issue for this type of analysis. Here, we introduce AMRomics, a minimalized microbial genomics pipeline that can work efficiently with big datasets. We use different bacterial data collections to compare AMRomics against competitive tools and show that our pipeline can generate similar results of interest but with better performance. The software is open source and is publicly available at https://github.com/amromics/amromics under an MIT license.

## Background

Whole genome sequencing (WGS) of bacterial isolates using the next-generation sequencing technology has progressively become the predominant method in clinical microbiology, public health surveillance, and disease control [1, 2, 3]. The ability to study the complete genetic information of a large number of bacterial genomes provides the potential to generate insights into the pathogenic genotype/phenotype relationships [4], pathogenic virulence transmissibility [5, 6] and antibiotic resistance tracking [7, 8]. The combination of genomics information and epidemiological data has been used frequently in disease control processes, such as rapid outbreak clustering investigation of the recent SARS-CoV-2 pandemic [9, 10] and evolutionary perspectives inference/prediction with regards to pathogenic diversification [11, 12]. The richness of current high-throughput genomic data has created a solid foundation to establish systematic studies for large cohorts of related genomes by applications of genome-wide methods such as cgMLST, phylogenetic, or pan-genomic analyses. WGS approaches can generate insightful data to discern knowledge about existing pathogenesis and assist in unraveling the characteristics of unknown ones [13, 14], which is critical in understanding and thus controlling disease outbreaks.

To meet the demand for analysis tools, a healthy number of computational pipelines have been developed to facilitate the analysis of microbial WGS data and to generate practical results of interest. Several have become well-established and widely used in the field, notably Nullarbor [15], Bactopia [16], and ASA^3^P [17]. The first-mentioned tool, Nullarbor, has been around as part of a standard process in public health microbial genomic procedure for the long haul, while the latter two are relatively up-to-date with comprehensive and wide-spectrum functionalities. However, these software pipelines usually require high-end computation infrastructures and take prohibitively long running times to analyze when collection sizes reach beyond thousands of genomes. Furthermore, while it is typical for laboratories to collect and sequence new samples over time, none of the existing pipelines can efficiently manage the growing collections where new samples are constantly added. In most cases, a large part of these pipelines need to be rerun every time new samples are added to the collection, resulting in additional high computation costs.

Here we introduce AMRomics, a lightweight open-source software for analyzing and managing large collections of bacterial genomes. This tool offers the ability to generate essential genomic results for individual samples, together with a population analysis that outperforms other methods. Thanks to its optimal design, the performance is significantly improved, making analyses of big collections of bacteria feasible on regular desktop computers with reasonable turn-around time. AMRomics project source code is available at https://github.com/amromics/amromics.git

### Workflow and Implementation

AMRomics is a software package that provides a comprehensive suite of genomics analyses of microbial collections in a simple and easy to use manner. It is designed to be performant and scalable to large genome collection with minimal hardware requirements without compromising the analysis results. To that end, we select the considered best practices tools in microbial genomics, and stitch them together via a well-structured workflow as described in the next section. For certain tasks in the workflow, AMRomics provides options for users to select among several alternative tools. The workflow is written in Python and is designed a modular and expandable application with the standardized data formats flowing between the tools in the workflow.

The software flexibly takes in input data in various formats including sequencing reads (with Illumina, Pacbio and Nanopore technologies), genome assembly, and genome annotations. It then performs assembly, genome annotation, MLST, virulome and resistome prediction, pangenome clustering, phylogenetic tree construction for each gene and core genes, and pan-SNPs analysis, all with a simple command line. AMRomics achieves this by building a pipeline consisting of the current best practice tools in bacterial genomics. It is also designed to be fast, efficient, and scalable to collections of thousands of isolates on a computer with modest hardware. Crucially, AMRomics supports the progressive analysis of a growing collection, where new samples can be added to an existing collection without the need to build the collection from scratch.

Functionally, the AMRomics pipeline can be split into 2 stages: single-sample analysis and pan-genome analysis as depicted in Figure 1. In the single-sample stage, every sample is processed based on the types of input data. Specifically, for Illumina sequencing data, fastp [18, 19] is employed for quality control, adaptor trimming, quality filtering and read pruning. The pre-processed reads are then subject to sequence assembly to generate a genome assembly. SKASE [20] is the method of choice for assemblying Illumina sequencing data for its speed, but the user can optionally choose to use SPAdes [21, 22] for slightly better N50 with the extra computation time. If long read data (Nanopore and Pacbio) are provided, the sample genome is assembled by Flye [23]. The assembly step can be skipped if the user provides the genome assembly in FASTA format as input to the pipeline. Next, the genome assembly is annotated with Prokka [24] unless the annotations are provided by the user. The gene sequences are extracted and stored in files at predefined locations. The genome sequence is also subject to multi-locus strain typing with pubMLST database of typing scheme for bacterial strains[25], antibiotic-resistant gene identification with AMRFinderPlus database[26], and virulent gene identification with the virulence factor database VFDB[27, 28]. All the results for single sample analysis are organized in a standard manner.

**Figure 1.**
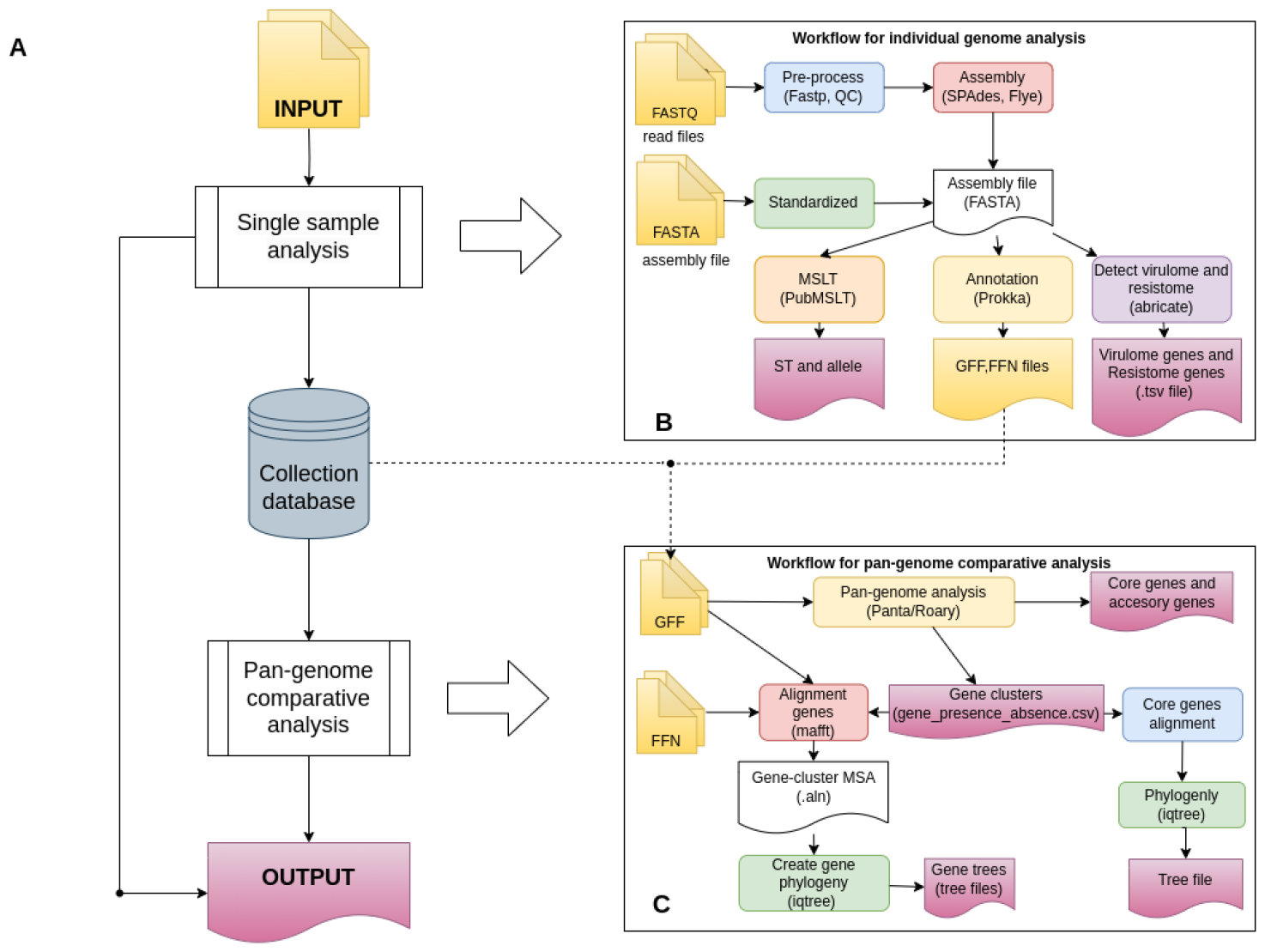
AMRomics workflow. **(A)** General framework of AMRomics including modules for single sample analysis and downstream comparative genomics for the collection. **(B)** Details of computational steps carried out first when adding individual isolates to the collection. **(C)** Details of downstream comparative analyses for the pan-genome collection.

In the second stage, AMRomics performs pan-genome comparative analysis of the genome collection. The annotations of all the genomes in GFF format are loaded into a pan-genome inference module for gene clustering. PanTA [29] is the method of choice for pan-genome construction for its speed and scalability, but users can optionally choose Roary [30] as the alternative. AMRomics then classifies gene clusters into core genes (genes clusters that present in at least 95% of genomes in the collection) and accessory genes. In addition, AMRomics identifies shell genes, which are those present in at least a certain number of genomes in the pangenome. The threshold for shell genes is defaulted at 25% but can be adjusted by users. AMRomics then performs multiple alignments (MSA) of all the identified shell genes using MAFFT [31]. The MSAs of these shell genes are then used to construct the phylogenetic trees of genes using FastTree 2 [32] or IQTree2 [33]. In addition, AMRomics builds the phylogeny of collection from the concatenation of the MSAs of all core genes using the chosen tree-building method.

AMRomics introduces pan-SNPs a novel concept to represent genetic variants of the samples in the collection. Existing variant analysis methods usually rely on a reference genome, and can only identify variants in the genes presenting in the reference genome. This severely limits the analysis to only a fraction of the genome of interests because of the high variability between isolates within a clade. In addition, it is often not possible to have a reference genome that can represent the whole collection, especially if the collection is diverse and growing. AMRomics addresses this by building the pan-reference genome for the collection from the representative genes of each of the gene clusters. It then identifies the variants of all genes in a cluster against the representative gene directly from the MSA. The variant profile of a sample is the concatenation of the variations of all its genes, reported in a VCF file.

The representative gene for a gene cluster is chosen such that comes from the earliest genome in the collection list. With this selection strategy, if the users have a preferred reference genome, they can place the reference genome first in the collection list so that genes from the reference genome will be the representatives in their perspective clusters. Moreover, as AMRomics supports continuously adding new samples into the collection, the selection strategy also ensures that the representative gene for a cluster does not change as the new samples are added into the collection, and that a new representative gene is added to the pan-reference genome only if a new cluster is created as the result of the collection expansion.

All results obtained from running AMRomics can be ultimately aggregated as the final output for reporting or customized visualizations for end users. Details of the third-party bioinformatic tools and databases used by AMRomics are listed in Supplementary Table 1 and 2.

## Results

### Comparison with other pipelines

To the best of our knowledge, at the time of writing, there are four existing open source software pipelines for end to end microbial genomics analysis, namely Nullarbor [15], TORMES [34], ASA^3^P [17] and Bactopia [16]. While AMRomics and these software tools share the overall functionalities, they differ in the underlying philosophies. Here, we present a high level discussion of AMRomics features and highlight the principles behind the design of AMRomics.

Overall, AMRomics and the existing tools support a wide variety of input formats except Nullarbor and TORMES which wre designed to run on Illumina paired-end reads only as per their specific public health routine. AMRomics and the more recent methods, ASA^3^P [17] and Bactopia accept raw reads from third-generation sequencing technology such as Oxford Nanopore Technology or PacBio long reads. A range of genomics analyses are included in all pipelines. They are common tasks for bacteria genomics such as sequence typing (MLST), AMR/virulence factor scanning, and genome annotation for an isolate. While all of the tools provide SNP analysis results, AMRomics outputs variants (in VCF files) by the core gene alignment from the pangenome analysis instead of snippy [35] core alignment as in other methods. Table 1 summarizes the key features across the software tools.

**Table 1:** Functional comparison between AMRomics and other bacterial genomics pipeline in use.

|  | AMRomics | Nullarbor | Bactopia | ASA<sup>3</sup>P |
| --- | --- | --- | --- | --- |
| Input | Long/short reads | Short reads | Long/short reads | Long/short reads |
|  | Assemblies |  | Assemblies | Assemblies |
|  | Annotated genomes |  |  | Annotated genomes |
| Output | Genotype | Genotype | Genotype | Genotype |
|  | AMR/Virulence | AMR/Virulence | AMR/Virulence | AMR/Virulence |
|  | Annotations | Annotations | Annotations | Annotations |
|  | SNP analysis | SNP analysis | SNP analysis | SNP analysis |
|  | Pangenomes | Pangenomes | Pangenomes | Pangenomes |
|  | Phylogenetics | Phylogenetics | Phylogenetics | Phylogenetics |
| Install | conda+pip | conda/docker | conda/docker | conda/docker |
| Progressive | Yes | No | Yes | No |

The primary principle of AMRomics is to extract the highest quality and most informative statistics from the input data. For example, AMRomics constructs the phylogeny tree of the collection using the multiple alignment of core genes. This provides a higher resolution of evolutionary information than SNPs information or the multiple alignment of 16S genes [36], the two techniques applied by the existing tools. In addition, AMRomics utilizes the population information to call variants across the pangenome instead of from a chosen reference genome and hence provides a bigger picture of genetic relations among the isolates in the collection. The users can still use one or more preferred reference genomes by placing the reference genomes at the top of the list.

AMRomics second and perhaps equally important design principle emphasizes on the on scallability of the software, aiming to be able to analyze large collections of genomes without the need to scale up hardware infrastructures. While AMRomics uses the same underlying core tools (*e*.*g*., BLAST+, SPAdes, SKASE, Flye, Prokka etc) as other pipelines, we chose to reimplement the helper and preprocessing modules such as Shovill and Dragonflye. In the process, we pay attention to the data structures to manage large amount of data flowing between steps of the pipeline. As a result, AMRomics is significantly faster and requires only a fraction of memory usage in comparison with its counterparts (shown in the following section). While speed is the paramount, AMRomics offers the flexibility for users to choose between alternatives to fit their need when there are more than one core algorithms for the same step (such as SPAdes and SKASE for assembling short reads, or FastTree and IQTREE for phylogenetic tree construction). AMRomics also takes advantages of progressive analysis; when new samples are added into an existing collection, AMRomics only performs the extra computation related to new the samples, instead of recomputing the scratch. This strategy offers a scalable solution practically suitable for analysis of the large growing collections of bacteria in the sequencing ages.

### Case study

We demonstrate the utility of AMRomics on a large and heterogeneous set of *Klebsiella pneumoniae* genomes collected from various public sources. In particular, we designed a case study that reflects a practical use case and highlights the ease of use, flexibility and scallability of AMRomics. The input data of the case study consisted of three batches of genome data. The first batch contained the sequencing data of 89 *K. pneumoniae* isolates from Patan Hospital in Kathmandu, Nepal between May and December 2012 [37]. These samples were multi-drug resistant isolates, in the form of Illumina paired-end short read data. While AMRomics did not require a reference genome for variant calling, we included in the batch four genome assemblies obtained from RefSeq (two in the genome assembly fasta format and two in annotation GFF format) for the other workflows to use as the reference. In the second batch, we included 11 samples that were collected from Hospital Universitario Ramon y Cajal, exhibiting Carbapenem resistance and harboring the pOXA-48 plasmid [38]. The input data for these 11 samples were Oxford Nanopore sequencing data. Finally, we included a third batch of 1000 samples; the genomes in the batch were previously assembled and annotated by NCBI PGAP, and they were in GFF format. The data in the case study are provided in the Supporting data.

Despite the commonalities among the analysis pipelines, having a direct comparison can be challenging due to the variations in the processing steps and the selection of different analysis tools within each pipeline. For simplicity, we used the default settings to run all existing pipelines that would cover essential analyses as shown in Table 1. We also with the best effort to use the parameters that the most compatible with AMRomics. We did not include TORMES in the comparison because of its resemblance to its predecessor, Nullarbor. The experiments were conducted on a cloud server with moderate performance, equipped with a 6-core 12-thread E-2286G processor, 32GB of RAM, and a 960GB SSD drive.

Table 2 shows the running time and resource consumption using the four pipelines. For the first batch, AMRomics took only 4.32 hours for performing single analysis on 89 samples, significantly faster than Bactopia and Nullarbor with 8.82 hours and 11.09 hours respectively even though the three pipelines use the similar underlying algorithms (SKASE for Illumina read assembly, Prokka for annotation and BLAST for virulome and resistome calling). This is likely due to better process management and parallelization implemented in AMRomics software. ASA^3^P took much longer, 22.24 hours as a result of using a slower assembly algorithm SPAdes that typically produced higher N50 quality assemblies. Of note, AMRomics, Bactopia and Nullarbor could optionally use SPAdes as the short read assembler. It is also worth noting that variation calling was part of single analysis in Bactopia, ASA^3^P and Nullarbor which also contributed to the extended single analysis time of these tools. AMRomics took under 1 hour for collection analyses, including pan-genome inference, multiple alignment of cloud genes, phylogenetic analyses of organisms and of every cloud gene, and SNP analysis. Nullarbor performed collection analysis in much shorter time, 0.19 hours albeit producing only pan-genome and core-gene phylogeny. Bactopia and ASA^3^P took significantly longer, 2.32 hours and 12.24 hours respectively. Taking together, AMRomics required less than half of the times of other tools for the whole pipeline. It also consumed only 3.44Gb of memory, comparing with 5.83Gb by Bactopia, 20.86Gb by ASA^3^P and 7.91Gb by Nullarbor.

**Table 2:** Running times and memory usages of the AMRomics, Bactopia, ASA^3^P and Nullarbor the case study.

|  | AMRomics |  | Bactopia |  | ASA<sup>3</sup>P |  | Nullarbor |  |
| --- | --- | --- | --- | --- | --- | --- | --- | --- |
|  | E. time<br>(Hrs) | M. mem<br>(Gb) | E. Time<br>(Hrs) | M. mem<br>(Gb) | E. time<br>(Hrs) | M. mem<br>(Gb) | E. time<br>(Hrs) | M. mem<br>(Gb) |
| **Batch 1** |  |  |  |  |  |  |  |  |
| Sample analysis | 4.32 | 3.44 | 8.82 | 5.83 | 22.24 | 6.71 | 11.09 | 7.91 |
| Collection analysis | 0.95 | 0.84 | 2.32 | 5.02 | 12.24 | 20.86 | 0.19 | 2.14 |
| Total | 5.27 | 3.44 | 11.14 | 5.83 | 34.48 | 20.86 | 11.28 | 7.91 |
| **Batch 2** |  |  |  |  |  |  |  |  |
| Sample analysis | 2.74 | 9.93 | 1.86 | 10.10 | 28.66 | 6.16 | - | - |
| Collection analysis | 0.94 | 0.89 | 3.99 | 5.74 | 26.10 | 28.87 | - | - |
| Total | 3.69 | 10.93 | 5.85 | 10.10 | 54.76 | 28.87 | - | - |
| **Batch 3** |  |  |  |  |  |  |  |  |
| Sample analysis | 4.17 | 1.22 | 67.72 | 4.21 | - | - | - | - |
| Collection analysis | 2.44 | 3.92 | - | - | - | - | - | - |
| Total | 6.61 | 3.92 | - | - | - | - | - | - |
| Accumulated | 9.06 | 10.93 |  |  |  |  |  |  |

The second batch consists of 11 Nanopore sequencing data, that was not supported by Nullarbor. ASA^3^P did not support progressive analysis hence all samples in the first batch and second batch had to be analyzed from scratch leading to a total of 54.76 hours. Bactopia took 1.86 hours for single analysis which was significantly shorter than AMRomics that took 2.74 hours though both tools used the same underlying assembly algorithm, Flye. Upon examining the runtimes, we noticed that Bactopia performed subsampling of sequencing reads to 50x resulting in the speed-up. AMRomics took less than one hour for collection analysis thanks to the use of progressive mode of its underlying pangenome method PanTA. On the other hand, Bactopia took 3.99 hours.

The genomes in the third batch were already annotated in GFF format. We did not run ASA^3^P on the third batch because of the excessive time required re-analyze the samples in the previous batches. Bactopia did not have the function to extract the annotations in the GFF files, and instead re-annotated the input genomes. In addition, Bactopia simulated sequencing reads from the assembled genomes, and mapped the simulated reads back to the reference to call SNPs. These steps, while could produce the intended analysis results, took 67.72 hours to analyze 1000 genomes. On the other hand, AMRomics reused the existing annotations from the input genomes, leading to substantially shorter single analysis running time, only 4.17 hours. Similarly, the pangenome analysis strategy employed by AMRomics reused the existing pangenome computation, requiring only 2.44 hours to add 1000 genomes into the existing pangenome. Bactopia ran pangenome analysis for more than 20 hours before crashing due to out of memory.

## DISCUSSION

We introduce AMRomics, a lightweight and scalable computational pipeline to analyze bacterial genomes and pan-genomes cost-effectively. The main focus of our method is to optimize the selected sub-modules for microbial genomic studies, especially comparative genomics, and most importantly to support progressive analysis for growing big data collections. AMRomics provides flexible input scenarios by supporting a wide range of data formats, such as different types of raw reads, assemblies, or annotated genomes for each sample. It can generate fundamental genomic properties sample-by-sample by carrying out routine analyses for bacteria isolates, and comparative genomics for the whole big collection *i*.*e*. pan-genome evaluation and the corresponding phylogenetic results.

## References

[1] Kwong JC, McCallum N, Sintchenko V, Howden BP (2015) Whole genome sequencing in clinical and public health microbiology. Pathology 47(3):199–210.

[2] Brown E, Dessai U, McGarry S, Gerner-Smidt P (2019) Use of whole-genome sequencing for food safety and public health in the united states. Foodborne pathogens and disease 16(7):441–450.

[3] Ferdinand AS et al. (2021) An implementation science approach to evaluating pathogen whole genome sequencing in public health. Genome Medicine 13:1–11.

[4] Karlsen ST, Rau MH, Sánchez BJ, Jensen K, Zeidan AA (2023) From genotype to phenotype: computational approaches for inferring microbial traits relevant to the food industry. FEMS Microbiology Reviews p. fuad030.

[5] Massey RC, Horsburgh MJ, Lina G, Höök M, Recker M (2006) The evolution and maintenance of virulence in staphylococcus aureus: a role for host-to-host transmission? Nature Reviews Microbiology 4(12):953–958.

[6] De la Fuente J et al. (2015) Comparative genomics of field isolates of mycobacterium bovis and m. caprae provides evidence for possible correlates with bacterial viability and virulence. PLOS Neglected Tropical Diseases 9(11):e0004232.

[7] Alghoribi MF, Balkhy HH, Woodford N, Ellington MJ (2018) The role of whole genome sequencing in monitoring antimicrobial resistance: A biosafety and public health priority in the arabian peninsula. Journal of Infection and Public Health 11(6):784–787.

[8] Hendriksen RS et al. (2019) Using genomics to track global antimicrobial resistance. Frontiers in public health 7:242.

[9] Petrone ME et al. (2022) Combining genomic and epidemiological data to compare the transmissibility of sars-cov-2 variants alpha and iota. Communications biology 5(1):439.

[10] Haanappel CP et al. (2023) Combining epidemiological data and whole genome sequencing to understand sars-cov-2 transmission dynamics in a large tertiary care hospital during the first covid-19 wave in the netherlands focusing on healthcare workers. Antimicrobial Resistance & Infection Control 12(1):1–12.

[11] Duault H, Durand B, Canini L (2022) Methods combining genomic and epidemiological data in the reconstruction of transmission trees: A systematic review. Pathogens 11(2):252.

[12] Khataei MM et al. (2022) A review of green solvent extraction techniques and their use in antibiotic residue analysis. Journal of Pharmaceutical and Biomedical Analysis 209:114487.

[13] Donkor ES (2013) Sequencing of bacterial genomes: principles and insights into pathogenesis and development of antibiotics. Genes 4(4):556–572.

[14] Li LM, Grassly NC, Fraser C (2014) Genomic analysis of emerging pathogens: methods, application and future trends. Genome biology 15(11):1–9.

[15] Seemann T GdSA (2018) Github https://github.com/tseemann/nullarbor.

[16] Petit III RA, Read TD (2020) Bactopia: a flexible pipeline for complete analysis of bacterial genomes. Msystems 5(4):10–1128.

[17] Schwengers O et al. (2020) ASA3P: an automatic and scalable pipeline for the assembly, annotation and higher-level analysis of closely related bacterial isolates. PLoS computational biology 16(3):e1007134.

[18] Chen S, Zhou Y, Chen Y, Gu J (2018) fastp: an ultra-fast all-in-one fastq preprocessor. Bioinformatics 34(17):i884–i890.

[19] Chen S (2023) Ultrafast one-pass fastq data preprocessing, quality control, and deduplication using fastp. iMeta p. e107.

[20] Souvorov A, Agarwala R, Lipman DJ (2018) SKESA: strategic k-mer extension for scrupulous assemblies. Genome Biology 19(1):153.

[21] Prjibelski AD et al. (2014) ExSPAnder: a universal repeat resolver for DNA fragment assembly. Bioinformatics 30(12):i293–i301.

[22] Vasilinetc I, Prjibelski AD, Gurevich A, Korobeynikov A, Pevzner PA (2015) Assembling short reads from jumping libraries with large insert sizes. Bioinformatics 31(20):3262–3268.

[23] Kolmogorov M, Yuan J, Lin Y, Pevzner PA (2019) Assembly of long, error-prone reads using repeat graphs. Nature biotechnology 37(5):540–546.

[24] Seemann T (2014) Prokka: rapid prokaryotic genome annotation. Bioinformatics 30(14):2068–2069.

[25] Jolley KA, Maiden MC (2010) BIGSdb: scalable analysis of bacterial genome variation at the population level. BMC bioinformatics 11:1–11.

[26] Feldgarden M et al. (2021) AMRFinderPlus and the reference gene catalog facilitate examination of the genomic links among antimicrobial resistance, stress response, and virulence. Scientific reports 11(1):1–9.

[27] Chen L et al. (2005) VFDB: a reference database for bacterial virulence factors. Nucleic acids research 33(suppl 1):D325–D328.

[28] Liu B, Zheng D, Zhou S, Chen L, Yang J (2022) Vfdb 2022: a general classification scheme for bacterial virulence factors. Nucleic acids research 50(D1):D912–D917.

[29] Le DQ et al. (2023) PanTA : An ultra-fast method for constructing large and growing microbial pangenomes. bioRxiv pp. 1–9.

[30] Page AJ et al. (2015) Roary: rapid large-scale prokaryote pan genome analysis. Bioinformatics 31(22):3691–3693.

[31] Katoh K, Asimenos G, Toh H (2009) Multiple alignment of dna sequences with mafft. Bioinformatics for DNA sequence analysis pp. 39–64.

[32] Price MN, Dehal PS, Arkin AP (2010) FastTree 2 – Approximately Maximum-Likelihood Trees for Large Alignments. PLoS ONE 5(3):e9490.

[33] Minh BQ et al. (2020) IQ-TREE 2: new models and efficient methods for phylogenetic inference in the genomic era. Molecular biology and evolution 37(5):1530–1534.

[34] Quijada NM, Rodríguez-Lázaro D, Eiros JM, Hernández M (2019) TORMES: an automated pipeline for whole bacterial genome analysis. Bioinformatics 35(21):4207–4212.

[35] Seeman T (2013) Github https://github.com/tseemann/snippy.

[36] Hassler HB et al. (2022) Phylogenies of the 16S rRNA gene and its hypervariable regions lack concordance with core genome phylogenies. Microbiome 10(1):104.

[37] Chung The H et al. (2015) A high-resolution genomic analysis of multidrug-resistant hospital outbreaks of klebsiella pneumoniae. EMBO molecular medicine 7(3):227–239.

[38] León-Sampedro R et al. (2021) Pervasive transmission of a carbapenem resistance plasmid in the gut microbiota of hospitalized patients. Nature microbiology 6(5):606–616.

