## Supplemental for "AMRomics: a scalable workflow to analyze large microbial genome collection"

**Supplementary Table 1.** List of bioinformatics tools used by AMRomics pipeline

| Tools | Source | Version | Function |
|---|---|---|---|
| | | | *Single sample analysis* |
| abricate | https://github.com/tseemann/abricate | 1.0.1 | Mass screening of contigs for antimicrobial and virulence genes[22] |
| blast | https://ftp.ncbi.nlm.nih.gov/blast/executables/blast+/LATEST/ | 2.13.0 | NCBI blast package for genome alignment[16] |
| fastp | https://github.com/OpenGene/fastp | 0.23.3 | Quick FASTQ reads trimming (adapters, length, quality)[4,5] |
| FastQC | https://github.com/s-andrews/FastQC | 0.11.9 | Quality control analysis for high throughput sequencing data[1] |
| Flye | https://github.com/fenderglass/Flye | 2.9-b1768 | *De novo* genome assembler for long-reads data[12,14] |
| mash | https://github.com/marbl/Mash | 2.3 | Fast genome size, (meta)genome distance estimation using MinHash[19] |
| mlst | https://github.com/tseemann/mlst | 2.19.0 | Scan contig files against PubMLST[9] typing schemes[21] |
| MultiQC | https://github.com/ewels/MultiQC | 1.10 | Aggregate results from bioinformatics analyses across many samples into a single report[7] |
| Prokka | https://github.com/tseemann/prokka | 1.14.6 | Rapid prokaryotic genome annotation[23] |
| samtools | https://github.com/samtools/samtools | 1.17 | Manipulate NGS alignment data[6] |
| seqtk | https://github.com/lh3/seqtk | 1.4-r122 | Processing, downsampling sequences data in FASTA/Q formats[13] |
| SPAdes | https://github.com/ablab/spades | 3.15.5 | *De novo* genome assembler for Illumina short-reads data[20,25] |
| | | | *Pan-genome comparative analysis* |
| MAFFT | https://github.com/GSLBiotech/mafft | 7.520 | Align multiple amino acid or nucleotide sequences[10,11] |
| msa2vcf | https://github.com/connor-lab/msa2vcf | | Turn a fasta-format MSA into one vcf per sequence.[?] |
| IQTREE | https://github.com/iqtree/iqtree2 | 2.2.2.3 | Build phylogenomics by maximum likelihood[17] |
| FastTree | http://www.microbesonline.org/fasttree/ | 2.1.11 | Build approximately-maximum-likelihood phylogenetic trees from large alignments of nucleotide[?] |
| Panta | https://github.com/amromics/panta | 1.0 | Fast bacterial pangenome analysis[?] |

**Supplementary Table 2.** List of bioinformatics databases used by AMRomics pipeline

| Database | Source | Description |
|---|---|---|
| PubMLST | https://pubmlst.org | MLST typing scheme for bacterial strains[9] |
| AMRFinderPlus | https://ftp.ncbi.nlm.nih.gov/pathogen/Antimicrobial_resistance/AMRFinderPlus/database/latest/ | Database of antimicrobial resistance genes and point mutations in the assemblies[8] |
| VFDB | http://www.mgc.ac.cn/VFs/Down/VFDB_setA_nt.fas.gz | Virulence factor database[3,15] |
| PlasmidFinder | https://bitbucket.org/genomicepidemiology/plasmidfinder_db | Database of origin of replicon sequences to identify plasmid[2] |
| INTEGRALL | http://integrall.bio.ua.pt/ | Database and search engine for integrons, integrases and gene cassette[18] |
| IntFinder | https://bitbucket.org/genomicepidemiology/intfinder_db | Integron database following INTEGRALL nomenclature[24] |

### References


1. S. Andrews. FastQC: A Quality Control Tool for High Throughput Sequence Data [Online]. 2010.
2. A. Carattoli, E. Zankari, A. García-Fernández, M. Voldby Larsen, O. Lund, L. Villa, F. Møller Aarestrup, and H. Hasman. In silico detection and typing of plasmids using plasmidfinder and plasmid multilocus sequence typing. *Antimicrobial agents and chemotherapy*, 58(7):3895–3903, 2014.
3. L. Chen, J. Yang, J. Yu, Z. Yao, L. Sun, Y. Shen, and Q. Jin. VFDB: a reference database for bacterial virulence factors. *Nucleic acids research*, 33(suppl_1):D325–D328, 2005.
4. S. Chen. Ultrafast one-pass fastq data preprocessing, quality control, and deduplication using fastp. *iMeta*, page e107, 2023.
5. S. Chen, Y. Zhou, Y. Chen, and J. Gu. fastp: an ultra-fast all-in-one fastq preprocessor. *Bioinformatics*, 34(17):i884–i890, 2018.
6. P. Danecek, J. K. Bonfield, J. Liddle, J. Marshall, V. Ohan, M. O. Pollard, A. Whitwham, T. Keane, S. A. McCarthy, R. M. Davies, and H. Li. Twelve years of SAMtools and BCFtools. *GigaScience*, 10(2), 02 2021. giab008.
7. P. Ewels, M. Magnusson, S. Lundin, and M. Käller. MultiQC: summarize analysis results for multiple tools and samples in a single report. *Bioinformatics*, 32(19):3047, 2016.
8. M. Feldgarden, V. Brover, N. Gonzalez-Escalona, J. G. Frye, J. Haendiges, D. H. Haft, M. Hoffmann, J. B. Pettengill, A. B. Prasad, G. E. Tillman, et al. AMRFinderPlus and the reference gene catalog facilitate examination of the genomic links among antimicrobial resistance, stress response, and virulence. *Scientific reports*, 11(1):1–9, 2021.
9. K. A. Jolley and M. C. Maiden. BIGSdb: scalable analysis of bacterial genome variation at the population level. *BMC bioinformatics*, 11:1–11, 2010.
10. K. Katoh, G. Asimenos, and H. Toh. Multiple alignment of dna sequences with mafft. *Bioinformatics for DNA sequence analysis*, pages 39–64, 2009.
11. K. Katoh and M. C. Frith. Adding unaligned sequences into an existing alignment using mafft and last. *Bioinformatics*, 28(23):3144–3146, 2012.
12. M. Kolmogorov, J. Yuan, Y. Lin, and P. A. Pevzner. Assembly of long, error-prone reads using repeat graphs. *Nature biotechnology*, 37(5):540–546, 2019.
13. H. Li. Github `https://github.com/lh3/seqtk`. 2011.
14. Y. Lin, J. Yuan, M. Kolmogorov, M. W. Shen, M. Chaisson, and P. A. Pevzner. Assembly of long error-prone reads using de bruijn graphs. *Proceedings of the National Academy of Sciences*, 113(52):E8396–E8405, 2016.
15. B. Liu, D. Zheng, S. Zhou, L. Chen, and J. Yang. Vfdb 2022: a general classification scheme for bacterial virulence factors. *Nucleic acids research*, 50(D1):D912–D917, 2022.
16. S. McGinnis and T. L. Madden. BLAST: at the core of a powerful and diverse set of sequence analysis tools. *Nucleic acids research*, 32(suppl_2):W20–W25, 2004.
17. B. Q. Minh, H. A. Schmidt, O. Chernomor, D. Schrempf, M. D. Woodhams, A. Von Haeseler, and R. Lanfear. IQ-TREE 2: new models and efficient methods for phylogenetic inference in the genomic era. *Molecular biology and evolution*, 37(5):1530–1534, 2020.
18. A. Moura, M. Soares, C. Pereira, N. Leitão, I. Henriques, and A. Correia. INTEGRALL: a database and search engine for integrons, integrases and gene cassettes. *Bioinformatics*, 25(8):1096–1098, 2009.
19. B. D. Ondov, T. J. Treangen, P. Melsted, A. B. Mallonee, N. H. Bergman, S. Koren, and A. M. Phillippy. Mash: fast genome and metagenome distance estimation using minhash. *Genome biology*, 17(1):1–14, 2016.
20. A. D. Prjibelski, I. Vasilinetc, A. Bankevich, A. Gurevich, T. Krivosheeva, S. Nurk, S. Pham, A. Korobeynikov, A. Lapidus, and P. A. Pevzner. ExSPAnder: a universal repeat resolver for DNA fragment assembly. *Bioinformatics*, 30(12):i293–i301, 06 2014.

**21.** T. Seeman. Github https://github.com/tseemann/mlst. 2014.

**22.** T. Seeman. Github https://github.com/tseemann/abricate. 2017.

**23.** T. Seemann. Prokka: rapid prokaryotic genome annotation. *Bioinformatics*, 30(14):2068–2069, 2014.

**24.** L. Torres-Elizalde, D. Ortega-Paredes, K. Loaiza, E. Fernández-Moreira, and M. Larrea-Álvarez. In silico detection of antimicrobial resistance integrons in salmonella enterica isolates from countries of the andean community. *Antibiotics*, 10(11):1388, 2021.

**25.** I. Vasilinetc, A. D. Prjibelski, A. Gurevich, A. Korobeynikov, and P. A. Pevzner. Assembling short reads from jumping libraries with large insert sizes. *Bioinformatics*, 31(20):3262–3268, 06 2015.
